# Germanium nanospheres for ultraresolution picotensiometry of kinesin motors

**DOI:** 10.1101/2020.06.18.159640

**Authors:** Swathi Sudhakar, Mohammad Kazem Abdosamadi, Tobias Jörg Jachowski, Michael Bugiel, Anita Jannasch, Erik Schäffer

## Abstract

Kinesin motors are essential for transport of cellular cargo along cytoskeletal microtubule filaments. How motors step, detach, and cooperate is still unclear. To dissect the molecular motion of kinesin-1, we have developed germanium nanospheres as ultraresolution optical trapping probes. We found that single motors took 4-nm-center-of-mass steps. Furthermore, motors never detached from microtubules under native hindering load conditions. Instead, motors slid on microtubules with microsecond-long, 8-nm steps and remained in this slip state before detaching or reengaging in directed motion. Surprisingly, reengagement and, thus, rescue of directed motion was prevailing. We argue that teams of motors may be synchronized through this slip state and rescues need to be accounted for to understand long-range transport.

**One Sentence Summary:** Optical trapping of high-refractive-index semiconductor nanoparticles shows how motors detach and walk with 4-nm steps.

---

Force spectroscopy on single molecular machines generating piconewton forces is often performed using optical tweezers (*1–3*). Since optical forces scale with the trapped particle volume, piconewton force measurements require micron-sized probes practically limiting the spatiotemporal resolution (*1, 2, 4, 5*). Fundamental limits arise due to Brownian motion that both molecular motors and trapping probes are subjected to (*5*). By temporal averaging over this motion, discrete motor steps of size *δ* and the time between steps—the dwell time *τ* — can be resolved. Such single-molecule measurements have provided unprecedented insight into essential mechanochemical processes of life (*1–3, 6*). However, many such processes cannot be measured at their native spatiotemporal resolution but only under conditions—for example, low nucleotide concentrations—at which the mechanochemistry is slowed down and might be different (*7*). For example, the benchmark, 3.4-Å–DNA-base-pair-sized steps of the RNA polymerase, naturally operating on a millisecond time scale, could only be resolved on a second time scale (*6*). The inherent trade-off between temporal and spatial precision and the resolution limit itself are quantified by the product 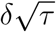 that has a constant value, with the lower limit hardly depending on the experiment (*2, 5*). Thus, this relation implies that detecting 8-nm steps of a kinesin motor on a millisecond time scale is as challenging as measuring Å-steps on a second time scale. Furthermore, apart from reducing linker compliance between probe and molecular machine, spatiotemporal resolution can only be significantly improved relative to the benchmark by the use of nanometer-sized optical trapping probes (*2, 5*). However, such probes for piconewton-force measurements do not exist.

Cytoskeletal motors like kinesins drive many essential cellular processes by coupling ATP hydrolysis to perform mechanical work (*8*). During an ATP hydrolysis cycle, kinesin motors advance by 8 nm along microtubules against forces of several piconewton via a rotational hand-over-hand mechanism (*7, 9*). While consensus develops on how kinesin motors work (*10, 11*) important details remain unclear. For example, it is controversial whether intermediate mechanical steps in the hydrolysis cycle exist and can support load (*12–17*). Furthermore, to enhance transport in crowded cells, kinesin motors work cooperatively in small teams (*18–20*). Key for team performance is how loads due to unsynchronized or opposing motors and obstacles affect transport distance (*18–21*). This distance and force generation are limited by motor detachment. However, how kinesins detach from microtubules is not known (*19, 21*).

To resolve how kinesin steps and detaches, we enhanced the spatiotemporal precision of optical tweezers by compensating the particle-volume-scaling of trapping forces in the Rayleigh regime with the use of highest infrared refractive index **ge**rmanium **n**anospheres as **t**rappable **o**ptical **p**robes (GeNTOPs). While various methods exist to make semiconductor nanoparticles (*22–26*), none provide water-stable, monodisperse, sufficiently large nanospheres for picotensiometry in adequate amounts. The synthesis that we developed derives from a solution-based method (*24*) and resulted in uniform GeNTOPs with a size of 72.0 ± 0.8 nm (mean ± standard error unless noted otherwise, *N* = 100) measured by transmission electron microscopy (TEM, Fig. 1A, see methods for details). To determine whether the spatiotemporal trapping precision of GeNTOPs was improved compared to commonly used microspheres, we trapped GeNTOPs in an ultrastable optical tweezers setup (*27*) (fig. S1) and calibrated them by a combined power spectral density–drag force method (*28, 29*) (fig. S2). The GeNTOP calibration showed that we achieved the optical-trap spring constant, the trap stiffness *κ*, necessary for kinesin picotensiometry (*7, 9, 12–14*). Also, for the used laser power, the trap stiffness quantitatively agreed with a Mie theory calculation based on the dielectric properties of germanium at the infrared trapping laser wavelength (see methods). Thus, GeNTOPs had indeed the expected very high refractive index of 4.4. In summary, because of the GeNTOPs’ high refractive index and nanometric size, spatial precision is significantly improved and the trap response time reduced by about an order of magnitude to *τ*_*trap*_ = (2*πf*_*c*_)^−1^ = *γ/κ* ≈ 10 μs, where *f*_*c*_ is the corner frequency and *γ* is the drag coefficient (fig. S2). By using a higher trap stiffness and/or smaller GeNTOPs—achieved through shorter reaction times—the response time can be reduced further.

**Fig. 1.**
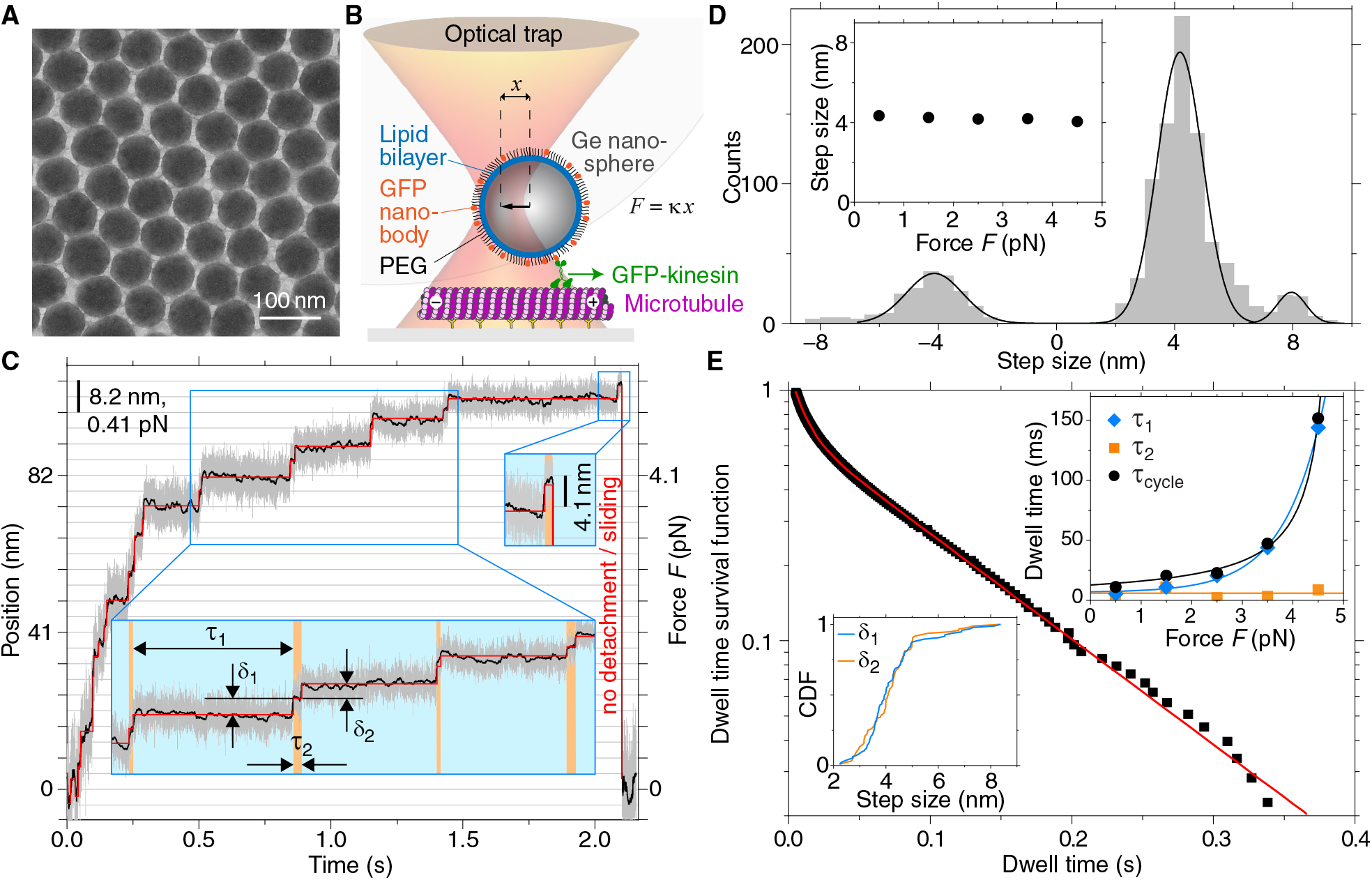
Ultraresolution kinesin traces employing optically trapped germanium nanospheres. **(A)**, TEM image of ≈70 nm germanium nanospheres (GeNTOPs). **(B)**, Schematic of a kinesin motor transporting a functionalized GeNTOP along a microtubule roughly drawn to scale including a section of a grey-shaded 0.59 μm diameter microsphere for comparison (the optical trap is too small; see text and methods for details). **(C)**, Time trace for a single-kinesin powered GeNTOP (100 kHz bandwidth, grey trace; filtered data, ≈100 Hz, black trace; detected steps, red line; see methods). Insets: magnified view of last and intermediate steps with definition of long and short dwell times *τ*_1_ (blue shaded) and *τ*_2_ (orange shaded) with corresponding step sizes *δ*_1_ and *δ*_2_, respectively. **(D)**, Step size histogram with a multi-Gaussian fit (line). Inset: Dominant step size versus force. **(E)**, Dwell time distribution of steps for *F* between 2–3 pN with fit (red line). Inset: Dwell times (symbols) with models (lines) versus force (top right, see methods); and cumulative distribution function (CDF) of alternating step sizes (bottom left).

To mimic *in vivo* vesicles while minimizing linker compliance and nonspecific interactions, we coated GeNTOPs with a PEGylated lipid bilayer functionalized with nanobodies that bound truncated, recombinant green-fluorescent-protein-(GFP)-tagged kinesin-1 motors hereafter called kinesin (Fig. 1B, fig. S3, see methods). The functionalization increased the GeN-TOP diameter to 93 ± 4 nm according to dynamic light scattering. This diameter corresponds to the average size of neuronal transport vesicles (*18*). Thus, dimensions and the force geometry when using GeNTOPs resemble native conditions inside cells. By using a low motor-to-GeNTOP ratio for further optical tweezers experiments, we ensured that only single kinesins transported GeNTOPs along microtubules with the expected speed and run length quantified by interference reflection microscopy (*30*) (fig. S3, see methods).

To dissect the kinesin gait, we trapped single-kinesin-functionalized GeNTOPs at physiological ATP concentrations, placed them on microtubules, and recorded the kinesin-powered GeNTOP displacement from the trap center (Fig. 1B). Based on this displacement *x* within the linear response of the GeNTOPs (inset fig. S2), the Hookean spring load of the optical tweezers corresponds to a force *F* = *κx*. In the exemplary trace of Fig. 1C (see more examples in fig. S4), motors slowed down with increasing hindering loads up to ≈5 pN. Also with increasing force, stepwise motion became more evident until GeNTOPs quickly returned to the trap center (in Fig. 1C at ≈2.1 s). To determine step sizes and dwell times, we used an efficient, automated filtering and step finding algorithm (see methods). Remarkably, instead of 8-nm steps (*9*), most forward-directed, center-of-mass steps were 4.12 ± 0.03 nm (center of Gaussian ± fit error) consistent with the size of a tubulin monomer. Because step size hardly depended on force (inset Fig. 1D, fig. S5), the combined linker-motor compliance was very low such that we could pool all steps together (Fig. 1D). There were only a few 8-nm forward and some 4-nm, but hardly any 8-nm, backward steps (table S1). Thus, our data directly shows that kinesin walks with 4-nm center-of-mass steps that can support load. Interestingly, for increasing forces, the step duration appeared to be alternating between a long and short dwell time that we denote by *τ*_1_ and *τ*_2_, respectively (Fig. 1C). Quantitatively, dwell time survival functions pooled from different force intervals were consistent with either a single exponential or sum of two exponentials with approximately equal amplitude for forces below or above 2 pN, respectively (Fig. 1E, table S1). Equal amplitudes imply that both type of dwells occurred equally often consistent with alternating steps having different properties. While the first dwell time *τ*_1_ depended on force, the second one, *τ*_2_, hardly depended on force (blue and orange lines in top right inset Fig. 1E, see methods). The sum of the two dwell times *τ*_*cycle*_ was consistent with a model based on the force-dependent speed of the motor (black circles and line in top right inset Fig. 1E, see methods) suggesting that each hydrolysis cycle is broken up into two mechanical substeps. Data recorded at low ATP concentrations (Fig. 2, see methods), show that only the first dwell time *τ*_1_ that depended on force also depended on ATP while *τ*_2_-values at low ATP hardly differed from the high-ATP values (table S1 and S2). Furthermore, for forces larger than 3 pN and physiological ATP concentrations, for which we could clearly assign alternating steps, the step size of alternating steps, always measured after the dwell, did not differ significantly (*δ*_1_ = 4.03 ± 0.06 nm, *N* = 97 and *δ*_2_ = 3.94 ± 0.06 nm, *N* = 88 for *τ*_1_ and *τ*_2_, respectively, bottom left inset Fig. 1E). However, we cannot rule out that the distributions consist of two closely spaced Gaussians with means that differ by the offset distance between neighboring protofilaments. Nevertheless, kinesin motors walked on average with 4-nm center-of-mass steps alternating in the force and ATP dependence of their dwell times.

**Fig. 2.**
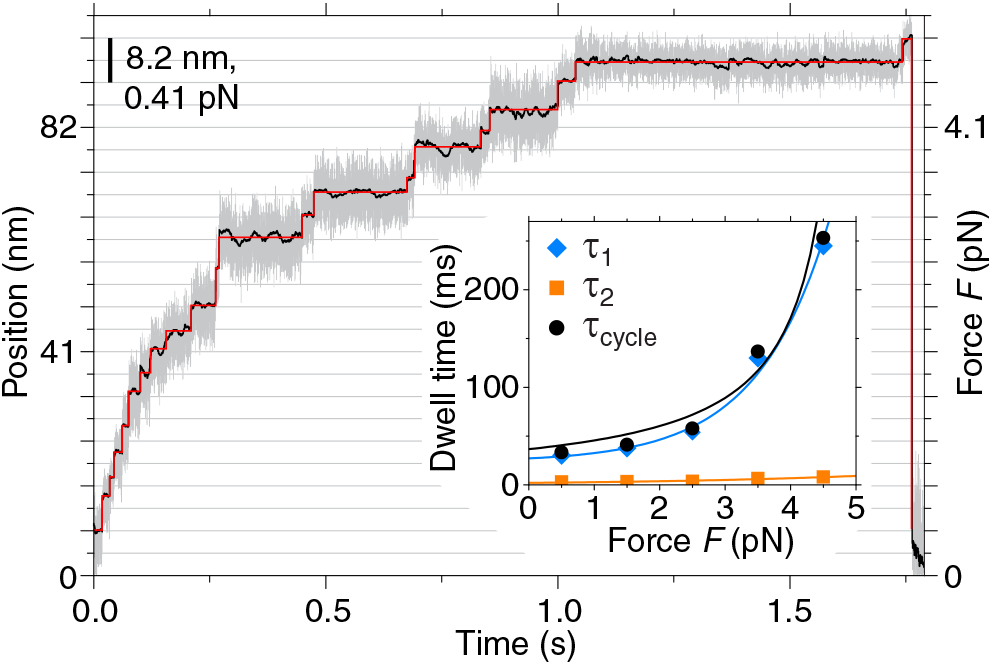
Low-ATP-concentration kinesin trace. Time trace for a single-kinesin powered GeNTOP at 10 μM ATP (100 kHz bandwidth, grey trace; filtered data, ≈100 Hz, black trace; detected steps, red line; see methods). Inset: Dwell times (symbols) with models (lines) versus force (see methods).

How and from which substep do motors detach? We noticed that in about 50 % of the motility events (*N* = 149), the last step before the GeNTOP quickly returned to the trap center was a short substep (Fig. 1C, fig. S4). For the subsequent fast backward motion, we expected an exponential relaxation with a time constant corresponding to the trap response time *τ*_*trap*_ in case of microtubule-motor detachment (*8*). However, while the backward motion directed along the microtubule axis could be fitted by an exponential relaxation (red line in Fig. 3A), the average time constant *τ*_॥_ = 295 ± 9 μs (*N* = 149)—and all individual ones without exception—was much larger than the trap response time. This discrepancy suggests that the kinesin still interacted with the microtubule and did not detach from it (Fig. 1C, Fig. 3A). To prevent microtubule interactions after the last step, we additionally pulled sideways on the kinesin-coated GeNTOP during motility events. With a load perpendicular to the microtubule axis, the relaxation time *τ*_⊥_ after the last step was only 30.7 ± 0.8 μs (*N* = 50) consistent with the expected trap relaxation time in the proximity of the surface (*29*) and true motor detachment (fig. S6). Close inspection of the relaxation traces along the microtubule (without sideward loads) revealed steps occurring on a microsecond time scale that were robustly detected by an unbiased change-point detection algorithm (*31*) (Fig. 3A and further examples in fig. S7 and S8, see methods). Individual steps were composed of an exponential relaxation with a time constant of 27 ± 3 μs (*N* = 20) consistent with the trap relaxation time *τ*_*trap*_ and had a step size of 7.2 ± 0.2 nm (*N* = 111, inset Fig. 3A)—close to the 8 nm repeat of the microtubule lattice—with a dwell time of 71 ± 4 μs (*N* = 124) averaged over all forces. Thus, we conclude that during fast backward motion, motors switched to a weakly bound slip state and remained in contact with the microtubule lattice. To determine whether motors truly detached from this weakly bound state or whether motors could switch back to a motility-competent state, we analyzed the time between subsequent motility events that we call restarting time (inset Fig. 3B). Intriguingly, also the restarting time survival function was well described by a sum of two exponentials having a time constant of 112 ± 1 ms and 4.1 ± 0.4 s, respectively (Fig. 3B). Two time constants imply that motility events started from two different states, possibly being *de novo* binding and the weakly bound state. The short restarting time constant that we measured is in excellent agreement with the one of a predicted weakly bound state prior to detachment of duration 131 ± 14 ms (*21*). Surprisingly, 82 ± 1 % of our events had this short restarting time constant suggesting that most motors did not detach but motility was rescued from the weakly bound state.

**Fig. 3.**
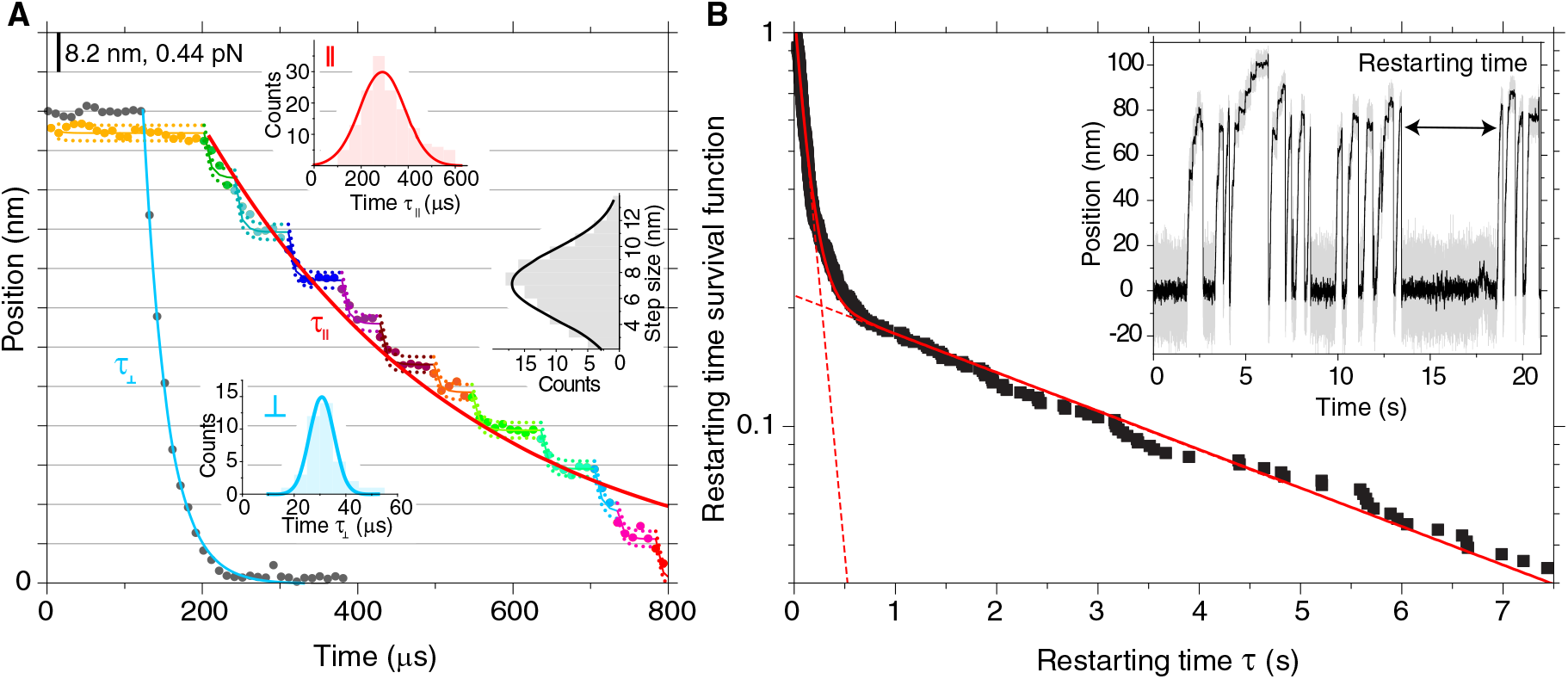
Ultrafast steps and motility rescue. **(A)**, Magnified time traces for a single-kinesin powered GeNTOP after the last step (grey and multicolored circles with or without sideward load, respectively, 100 kHz bandwidth) with single exponential fits (blue and red line for motion perpendicular (⊥) and parallel (॥) to the microtubule axis, respectively). Multicolored lines correspond to states detected by a change-point algorithm (*31*) (dotted line 95 % confidence interval). Inset: histograms with Gaussian fits (solid lines) of relaxation time constants *τ*_॥_ and *τ*_⊥_ (same color code as single exponential fits) and step size for detected states. **(B)**, Restarting-time distribution (squares) fitted with a sum of two exponentials (line with dashed line extrapolation, *N* = 550). Inset: Illustration of the restarting time between consecutive kinesin motility events.

Our data is consistent with a model for kinesin stepping that splits up the hydrolysis cycle into two mechanical substeps. In between the substeps, the motor can branch off from the normal hydrolysis pathway and switch to a weakly bound diffusive or sliding state prior to detachment or rescue of motility (Fig. 4). Overall, our model builds on and expands previous models (*7, 10, 11, 16, 32*). Initially, motor heads are bound to the microtubule with ADP and inorganic phosphate (P_*i*_) in the rear head and no nucleotide in the front one. With ATP binding to the front head and P_*i*_ release from the rear one, the rear neck linker is undocked and the front one docked. This process triggers the first 4-nm, ATP-dependent center-of-mass substep (Substep *τ*_1_(*F*,ATP) in Fig. 4). Since load is acting on the front neck linker during docking, it may explain that the dwell time of this step also depends on force. Upon ATP hydrolysis in the front head and ADP release from the rear one, the hydrolysis cycle is completed with a second 4-nm substep (Substep *τ*_2_ in Fig. 4). Since ATP is already bound, this substep does not depend on the ATP concentration. Also, because load is mainly acting on the rear head through the docked neck linker and the front head is free to perform a diffusive search with an undocked neck linker, may explain why the dwell time of this step hardly depends on force (inset Fig. 1E and Fig. 2). Based on previous (*7*) and our current data, we suggest that heads always remain weakly bound to the microtubule lattice likely due to electrostatic interactions, for example, with the negatively charge E-hooks of tubulin. If P_*i*_ is released from the front head directly after ATP hydrolysis and before ADP is released from the rear head, both heads enter a weakly bound, diffusive ADP state interrupting the normal hydrolysis cycle (red box in Fig. 4). Load will bias such a diffusive state—as observed for the fast backward movement after the last kinesin step when stalling—resulting in stepwise sliding motion opposed by hydrodynamic drag and protein friction (*33*). The measured step size of these fast, sliding steps close to 8 nm suggests that the motor heads interact primarily with the canonic kinesin-microtubule binding site. While we hardly observe 8-nm backward steps, we observed some short slip events (fig. S8). With a different force geometry and large microspheres that cause a large drag, such events may correspond to previously observed backward steps (*14*). Protein friction allows us to estimate the diffusive step dwell time during the fast sliding motion. Based on the time constant for the fast movement back to the trap center *τ*_‖_ = *τ*_*trap*_ + *γ_proteinfriction_/κ*, the force-averaged friction coefficient due to friction between the motor and its track is *γ*_*proteinfriction*_ ≈ 15 nN s/m and the corresponding diffusion coefficient according to the Einstein relation is *D* = *k_B_T*/*γ*_*proteinfriction*_ ≈ 0.3 μm^2^/s, where *k*_*B*_ is the Boltzmann constant and *T* the absolute temperature. Furthermore, if we model the backward movement by a biased one-dimensional random walk with a step size of *δ* = 8 nm, the expected average step time is *τ* ≈ *δ*^2^*/*(2*D*) ≈ 70 μs. This time constant is in excellent agreement with the directly measured dwell time during the fast backward sliding motion (Fig. 3A) and supports the notion of a biased weakly bound slip state prior to detachment or rescue (*21*). Unexpectedly, in only roughly 20 % of events, motors did truly detach, but in 80 % of the cases ADP must have dissociated from one of the heads rescuing directed motion (green box in Fig. 4). We expect that motors also switch to this diffusive state when no load is applied, suggesting that overall run lengths of motors are concatenations of processive runs interrupted by short diffusive periods (*34, 35*).

**Fig. 4.**
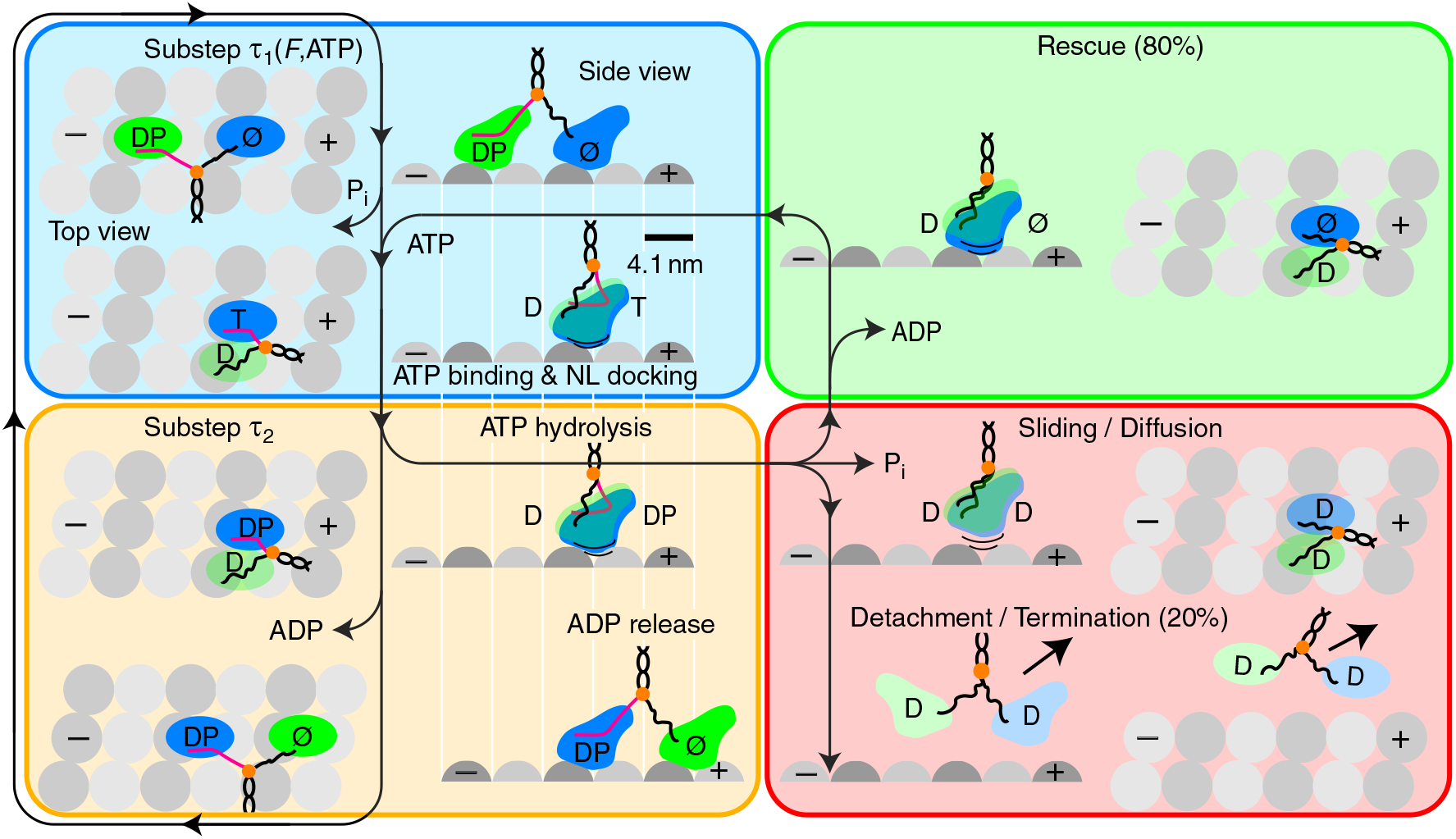
Hydrolysis cycle with detachment and rescue. Top and side view of kinesin with two identical heads (blue and green) stepping along a microtubule (grey spheres mark tubulin monomers). The hydrolysis cycle is divided into a force-dependent (blue box) and hardly force-dependent (orange box) substep with dwell times *τ*_1_(*F*,ATP) and *τ*_2_, respectively. Between these substeps, motors may switch to a weakly bound sliding or diffusive state from which motors either detach (red box) or motility is rescued (green box). The center of mass is indicated by an orange circle, a docked neck linker (NL) marked by a magenta line, weak binding by lines underneath the heads, and nucleotide states by T: ATP, D: ADP, P_*i*_: inorganic phosphate, and Ø: nucleotide free.

In general, widely available, size-controllable high-refractive index GeNTOPs will enable other applications due to having the highest infrared refractive index of common materials and being a semiconductor. Germanium nanospheres are a lower-toxicity alternative to compound semiconductor nanoparticles (*22, 24*), optimal for bioimaging and sensing at wavelengths biological tissues are transparent (*23*), promising for nanophotonics and optoelectronics (*25, 26*), and may enhance energy harvesting and storage (*36*). As for optical trapping and relative to the benchmark (*6*), the spatiotemporal resolution 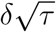 of the fast 8-nm steps on microsecond time scales, is an improvement by a factor of about 4.5× and 20× with respect to spatial and temporal resolution (fig. S9). Thus, GeNTOPs do allow to observe molecular machines at their native spatiotemporal resolution. In our case, the dwell time of the weakly bound state cannot be slowed down by reducing nucleotide concentrations because nucleotides likely did not exchange during sliding. For kinesins, the detachment and rescue state allows motors to slide back to their team during transport with direct reengagement in motility. This process provides a route for load distribution and motor synchronization enhancing transport. Therefore, for a better understanding of long-range transport in crowded cells (*19*) and of other essential cellular functions of kinesins, the sliding and rescue processes need to be accounted for.

## Acknowledgments

We thank Ulrich Rothbauer (NMI, Reutlingen, Germany) for providing the anti-GFP nanobody, Andreas Schnepf for the use of the Zetasizer, and Mohammed Mahamdeh, Joe Howard, Martin Oettel, and Carolina Carrasco for comments on the manuscript.

## Author contributions

S.S., and E.S. designed research; S.S. performed all experiments; S.S., M.K.A., A.J. and E.S. analyzed data; M.K.A and T.J.J. provided data analysis software; T.J.J. developed the Python package stepfinder; M.B. and A.J. developed protocols, controlled statistics, and provided advice; and S.S., and E.S. wrote the paper. All authors commented on the manuscript.

## Data and materials availability

All data are available in the main text and the supplementary materials.

## Supplementary materials

Materials and Methods

Figs. S1 to S9

Tables S1 to S2

References *(37–70)*

## References and Notes

1. K. Svoboda, S. M. Block, Annu. Rev. Biophys. Biomol. Struct. 23, 247 (1994).

2. J. R. Moffitt, Y. R. Chemla, S. B. Smith, C. Bustamante, Annu. Rev. Biochem. 77, 205 (2008).

3. A. Gennerich, Optical Tweezers, Methods Mol. Biol. (Springer, New York, 2017).

4. A. Ashkin, J. M. Dziedzic, J. E. Bjorkholm, S. Chu, Opt. Lett. 11, 288 (1986).

5. F. Gittes, C. F. Schmidt, Methods Cell Biol., M. P. Sheetz, ed. (Academic Press, 1997), vol. 55, pp. 129–156.

6. E. A. Abbondanzieri, W. J. Greenleaf, J. W. Shaevitz, R. Landick, S. M. Block, Nature 438, 460 (2005).

7. A. Ramaiya, B. Roy, M. Bugiel, E. Schäffer, Proc. Natl. Acad. Sci. USA 114, 10894 (2017).

8. J. Howard, Mechanics of motor proteins and the cytoskeleton (Sinauer Associates, Sunderland, MA, 2001).

9. K. Svoboda, C. F. Schmidt, B. J. Schnapp, S. M. Block, Nature 365, 721 (1993).

10. W. O. Hancock, Biophys. J. 110, 1216 (2016).

11. R. A. Cross, Biopolymers 105, 476 (2016).

12. C. M. Coppin, J. T. Finer, J. A. Spudich, R. D. Vale, Proc. Natl. Acad. Sci. U. S. A. 93, 1913 (1996).

13. M. Nishiyama, E. Muto, Y. Inoue, T. Yanagida, H. Higuchi, Nat. Cell Biol. 3, 425 (2001).

14. N. J. Carter, R. A. Cross, Nature 435, 308 (2005).

15. S. M. Block, Biophys. J. 92, 2986 (2007).

16. K. J. Mickolajczyk, et al., Proc. Natl. Acad. Sci. USA 112, E7186 (2015).

17. H. Isojima, R. Iino, Y. Niitani, H. Noji, M. Tomishige, Nat. Chem. Biol. 12, 290 (2016).

18. A. G. Hendricks, et al., Curr. Biol. 20, 697 (2010).

19. Q. Feng, K. J. Mickolajczyk, G.-y. Chen, W. O. Hancock, Biophys. J. 114, 400 (2018).

20. K. I. Schimert, B. G. Budaitis, D. N. Reinemann, M. J. Lang, K. J. Verhey, Proc. Natl. Acad. Sci. USA 116, 6152 (2019).

21. H. Khataee, J. Howard, Phys. Rev. Lett. 122, 188101 (2019).

22. J. Fan, P. K. Chu, Small 6, 2080 (2010).

23. D. D. Vaughn II, R. E. Schaak, Chem. Soc. Rev. 42, 2861 (2013).

24. Y. J. Guo, et al., Chem. Asian J. 9, 2272 (2014).

25. A. I. Kuznetsov, A. E. Miroshnichenko, M. L. Brongersma, Y. S. Kivshar, B. Luk’yanchuk, Science 354, aag2472 (2016).

26. A. Krasnok, M. Caldarola, N. Bonod, A. Alú, Adv. Opt. Mater. 6, 1701094 (2018).

27. M. Mahamdeh, E. Schäffer, Opt. Express 17, 17190 (2009).

28. S. F. Tolic-Nørrelykke, E. Schäffer, J. Howard, F. S. Pavone, F. Jülicher, Rev. Sci. Instrum. 77, 103101 (2006).

29. E. Schäffer, S. F. Nørrelykke, J. Howard, Langmuir 23, 3654 (2007).

30. S. Simmert, M. Abdosamadi, G. Hermsdorf, E. Schäffer, Opt. Express 26, 1437 (2018).

31. P. A. Wiggins, Biophys. J. 109, 346 (2015).

32. J. O. L. Andreasson, et al., eLife 21, 1 (2015).

33. V. Bormuth, V. Varga, J. Howard, E. Schäffer, Science 325, 870 (2009).

34. A. Jannasch, V. Bormuth, M. Storch, J. Howard, E. Schäffer, Biophys. J. 104, 2456 (2013).

35. M. Chugh, et al., Biophys. J. 115, 375 (2018).

36. T. H. Kim, H. K. Song, S. Kim, Nanotechnology 30, 275603 (2019).

