## Supplementary Materials for "Germanium nanospheres for ultraresolution picotensiometry of kinesin motors"

### 1. Materials and Methods

#### 1.1. Synthesis of germanium nanospheres (GeNTOPs)

The germanium nanospheres were synthesized in an aqueous solution advancing a method of Guo *et al.* (*37*). As substrate, 17.0 mg of germanium oxide ($GeO_2$) and 96.0 mg of quercetin, acting as a stabilizing agent, were dissolved in 10 ml of a 0.15 M sodium hydroxide solution each and then mixed together while stirring for 10 min and adjusting the pH to 8.8 via titration with 37 % HCl (Solution A). Subsequently, 29.5 mg of sodium borohydride ($NaBH_4$, reducing agent) was dissolved as quickly as possible in 3 ml of 4 °C-cold water and stored in a refrigerator at 4 °C (Solution B). Then, Solution A was stirred continuously in a preheated oil bath at 60 °C for 10 min and Solution B was added dropwise. The reaction was stopped after 5 h and GeNTOPs washed thrice thoroughly with water by centrifuging the sample at 13,000 rpm. All chemicals were purchased from Sigma Aldrich and used without further purification unless noted otherwise. Purified Type 1 water was used for all experiments (18.2 MΩ cm, Nanopure System MilliQ reference with Q-POD and Biopak filter). The size characterization analysis was done using a TEM-Jeol 1400 plus transmission electron microscope. About 10 µl of the GeNTOP solution was sonicated and subsequently 5 µl spotted on a TEM grid. Dynamic light scattering resulted in a diameter of $74 \pm 3$ nm consistent with the value obtained by TEM.

#### 1.2. Lipid-bilayer functionalization of GeNTOPs

After the synthesis, GeNTOPs were coated with a lipid bilayer using established methods (*38*–*40*). Briefly, 1,2-dimyristoyl-sn-glycero-3-phosphocholine (DMPC, Avanti Polar Lipids, Inc.) and 1,2-distearoyl-sn-glycero-3-phosphoethanolamine-N-[carboxy(polyethylene glycol)-2000] (DSPE-COOH, Avanti Polar Lipids, Inc.) were dissolved in chloroform (10 mg/ml). Aliquots of a 4:1 molar ratio mixture of these lipids were dried overnight in a desiccator at 50 mbar and stored at −20 °C. The dried lipid mixture was hydrated by adding 1 ml of 80 °C warm buffer (10 mM 4-(2-hydroxyethyl)-1-piperazineethanesulfonic acid (HEPES), 150 mM NaCl, pH 7.4) resulting in a final total lipid concentration of about 0.5 mM. To form multilamellar vesicles (MLVs), the solution was mixed thoroughly by pipetting and vortexed for 2 min. Subsequently, small unilamellar vesicles (SUVs) were formed by sonicating the MLV mixture for 30 min at 80 °C. The sonicated solution was centrifuged at 12,000 rpm for 15 min and SUVs collected from the supernatant. Then, equal volumes of GeNTOP and SUV solutions were mixed. To induce fusion of the liposomes onto the GeNTOPs, $CaCl_2$ was added to the mixture (3 mM final concentration) that was incubated for 45 min at 80 °C in a thermomixer using a shaking speed of 600 rpm. The membrane-coated GeNTOPs were washed thrice in three different buffers, first with Buffer 1 (25 mM HEPES, 200 mM NaCl, 1 mM tris(2-carboxyethyl)phosphine (TCEP), pH 7.4, 5 mM EDTA) followed by washing them in Buffer 2 (25 mM HEPES, pH 7.4, 100 mM NaCl, 0.25 mM $CaCl_2$) and then in Buffer 3 (25 mM HEPES, pH 7.4, 25 mM NaCl, 1 mM TCEP, 0.25 mM $CaCl_2$). After each wash, GeNTOPs were collected by spinning the sample at 13,000 rpm for 15 min and gently resuspending them. After the last resuspension step, GeNTOPs were lyophilized and kept at 4 °C for later use. For membrane visualization, 10 µl of 2 µM DiI lipophilic dye was added when hydrating the lipid mixture used to coat GeNTOPs. As a control, 100 µl of uncoated GeNTOPs, was mixed with 10 µl of 2 µM DiI lipophilic dye and incubated for 45 min. After incubation, these GeNTOPs were washed thrice with water and suspended in 100 µl water. Both the coated GeNTOPs with the membrane dye and control GeNTOPs were imaged by a Leica TCS SP8 confocal microscope with an excitation wavelength of 565 nm. No fluorescence was observed for the control.

### 1.3. GeNTOP PEGylation and nanobody coupling

For kinesin experiments, we PEGylated GeNTOPs and covalently bound GFP nanobodies to them as described previously (*41*) with some modifications. About 0.1 g of lyophylized GeNTOPs were dissolved in 1 ml water. From this stock, 25 µl were washed twice with 975 µl of 2-(N-morpholino)ethanesulfonic acid (MES) buffer (50 mM, pH = 6.0) by centrifuging GeNTOPs at 13,000 rpm for 15 min. Before each wash cycle, GeNTOPs were vortexed and sonicated in a bath sonicator for 15 s. Then, GeNTOPs were resuspended in 250 µl MES buffer. After washing, GeNTOPs were vortexed and sonicated for 180 s. Then, 16.4 mg of 1-(3-(dimethylamino)propyl)-3-ethylcarbodiimide hydrochloride (EDC) and 8.3 mg of N-hydroxysulfosuccinimide sodium (NHS) were dissolved in 100 µl of MES buffer. From the prepared solution, 9 µl of NHS and 15.8 µl of EDC were added to the resuspended GeNTOPs and the solution was mixed in a thermomixer for 15 min at 37 °C. Then, GeNTOPs were washed twice with 500 µl of MES buffer, resuspended in 240 µl of PBS-T (phosphate buffer saline supplemented with 0.1 % Tween 20), and vortexed and sonicated for 90 s. Subsequently, GFP-nanobodies (*42*) (13 kDa, gift of Ulrich Rothbauer, NMI, Reutlingen, Germany) and 2 kDa $\alpha$-methoxy-$\omega$-amino PEG (Rapp Polymere, Tübingen, Germany) in a molar ratio of 1:1000 were coupled covalently to the GeNTOPs by incubating them in a thermomixer for 1 h at 600 rpm and 37 °C. Afterwards, GeNTOPs were washed five times with PBS-T and stored at 4 °C.

### 1.4. Sample preparation and assay

Experiments were performed in flow cells that were constructed using silanized, hydrophobic glass cover slips and parafilm as described before (*43, 44*) but chlorotrimethylsilane (Merck Millipore, Burlington, MA) was used to render surfaces hydrophobic. Truncated kinesin1-eGFP-6xHis (rk430) was purified as described previously (*44, 45*). Taxol-stabilized microtubules, sometimes additionally 10% rhodamine-labeled, were prepared as described previously (*46*). Flow channels were washed with PEM buffer (80 mM 1,4-piperazinediethanesulfonic acid (PIPES), 1 mM EGTA, 1 mM $MgCl_2$, adjusted with KOH to pH 6.9), filled and incubated successively with anti $\beta$-tubulin I (monoclonal antibody SAP.4G5 from Sigma in PEM) for 15–20 min, Pluronic F-127 (1 % in PEM) for 20 min, and microtubules in PEM for 15 min. Kinesin with a stock concentration of 12.1 mg/ml was diluted 1000× in motility buffer (PEM with 0.16 mg/ml casein, 1 mM or 10 µM ATP and an anti-fade cocktail [20 mM D-glucose, 0.02 mg/ml glucose oxidase, 0.008 mg/ml catalase and 10 mM dithiothreitol]). Then 4 µl of the kinesin solution was mixed with 96 µl of 10× diluted functionalized GeNTOPs and incubated for 10 min. About, 20 µl of this GeNTOP-motility solution was flown into the channel for single-molecule force measurements. To rule out artifacts from angled motion in the optical trap (*47*), only microtubules aligned with the flow cell channel direction and perpendicular to the laser polarization (*48*) were chosen for experiments.

### 1.5. Optical tweezers setup and calibration

Measurements were performed in our ultraprecision optical tweezers setup (*43,49*). Briefly, the setup has near-Å resolution in surface-coupled assays (fig. S1) and is equipped with a millikelvin precision temperature control set to 29.500 °C (*49*). Signals of a 1064 nm trapping laser were recorded with 100 kHz by back focal plane detection. The optical trap was calibrated by a combined power spectral density–drag force method (*43, 50*). The average trap stiffness used for experiments was about 0.05 pN/ nm. For the power spectra in fig. S2, the trap stiffness was 0.0552 ± 0.0005 pN/nm and 0.0561 ± 0.0005 pN/nm recorded at 2 µm and 5 µm distance from the surface using about 600 mW and 6.5 mW of laser power in the focus for the GeNTOP and polystyrene microsphere, respectively. Both trap stiffness values quantitatively agreed with Mie theory calculations for our setup (*48, 51, 52*) using a refractive index of 4.4 + 0.11$i$ (*53*) for the GeNTOPs. Due to absorption, we measured a temperature increase for the GeNTOPs at 600 mW trapping power of about 7 K above the flow cell temperature 500 nm away from the surface using our calibration method (*54*). This temperature increase is slightly more than what is expected for heating due to the trapping laser alone (*55*). Since the surface acts as a heat sink (*55*), we expect that during kinesin experiments heating was less. We did not notice any significant changes due to temperature, e.g. in motor speed or force generation, compared to when using polystyrene microspheres with the same trap stiffness.

### 1.6. Step detection and data processing

For step detection and filtering, data was processed using an optimized, automated step finding algorithm (*56*) based on a modified forward-and-backward filter from Chung & Kennedy that we implemented in Python (*44, 56–58*). The filter works very efficiently in particular, for large data sets consisting of millions of data points. For sufficiently large data sets, the algorithm automatically finds the optimal window length for filtering and step detection according to the following idea: if we smooth the signal with different window lengths, the standard deviation of the smoothed signal decreases with increasing window length as long as the window length is shorter than the dwell time of the steps. As soon as the window includes steps, i.e. is comparable to the dwell time of the steps, the standard deviation increases again. The window length with the lowest standard deviation is used as a proxy for the optimal window size that we empirically chose to be 4/5 of the latter window length. To filter the data while preserving steps, the optimal window size is used to calculate the variance-weighted mean of the forward and backward window corresponding to the filtered data point. For our data, we used a window size of 4.8 ms. For step detection during the fast backward motion, we used the unbiased "Steppi" algorithm (*59*). In selected traces (fig. S7), the algorithm detected steps corresponding to single exponential relaxations with a time constant consistent with the trap response time. To robustly detect sliding steps in many traces, we fixed the relaxation time constant to the expected and exemplarily verified one. To account for the different trap response times in the different directions parallel and perpendicular to the microtubule axis and assuming that the hydrodynamic drag coefficient is the same in both directions (*43*), we chose a relaxation time of $\tau_\perp\kappa_\perp/\kappa_\parallel = 25\,\mu s$, where $\kappa_\parallel$ and $\kappa_\perp$ are the trap stiffness in the direction of the microtubule axis and perpendicular to it, respectively, and $\tau_\perp$ is the experimentally measured value (Fig. 3A). The average trap stiffness of $\kappa_\parallel$ and $\kappa_\perp$ was $0.051 \pm 0.001$ pN/nm ($N = 149$) and $0.041 \pm 0.001$ pN/nm ($N = 50$). To apply sideward loads during a motility event, we manually displaced the sample 50 nm in a direction perpendicular to the microtubule axis and relative to the stationary optical trap using a piezo-translation stage resulting in sideward loads of about 2 pN. For the last short step, we measured a dwell time of $58 \pm 12$ ms ($N = 74$) longer than the average $\tau_2$ value at that force indicating that the small increase of $\tau_2$ with force promotes the switching to the diffusive state. Speeds as a function of force are based on linear fits to trace segments in the respective force intervals, where automatic threshold detection of force was based on the filtered data. The speed (table S1 and S2) was well described by a linear force-velocity relation with zero-load speed $\upsilon_0 = 0.64 \pm 0.02\,\mu$m/s and $0.22 \pm 0.02\,\mu$m/s and stall force $F_s = 4.92 \pm 0.03$ pN and $5.1 \pm 0.7$ pN for high and low ATP concentrations, respectively. Based on this relation and fitted parameters, the total dwell time for a hydrolysis cycle is $\tau_{\text{cycle}} = (2\delta)/(\upsilon_0(1 - F/F_s))$ (black line in top right inset Fig. 1E and inset Fig. 2), where we used $\delta = 4.1$ nm. The force dependence of the substeps was modeled by $\tau(F) = \tau_0 \exp(Fx^\ddagger/(k_\text{B}T)) + \tau_{\text{const}}$, where for 1 mM ATP and the long dwell time $\tau_1$ the zero-force dwell time $\tau_0$ was $0.5 \pm 0.2$ ms, the distance to the transition state $x^\ddagger$ was $5.3 \pm 0.4$ nm, and the constant $\tau_{\text{const}}$ was $7 \pm 2$ ms (blue line in top right inset Fig. 1E). For 1 mM ATP and the short dwell time $\tau_2$, the data was best modeled by a constant value of $6.0 \pm 1.6$ ms (orange line in top right inset Fig. 1E). Note that for $F < 2$ pN, a single exponential modeled the data best and we used the same value for $\tau_1$ and $\tau_2$. For 10 µM ATP, the zero-force dwell time $\tau_0$ was $4 \pm 2$ ms and $2.1 \pm 0.4$ ms, the distance to the transition state $x^\ddagger$ was $3.8 \pm 0.6$ nm and $1.2 \pm 0.3$ nm, and the offset $\tau_{\text{const}}$ was $24 \pm 6$ ms and zero for $\tau_1$ and $\tau_2$, respectively (blue and orange line in inset Fig. 2).

#### 1.7. Single-molecule conditions

We measured the fraction of motile GeNTOPs $p_m \pm (p_m(1-p_m)/N)^{1/2}$ (mean ± error bar) by trapping GeNTOPs incubated with different concentrations of kinesin motors and placing them on microtubules to await motility (*41, 60*). The probability that a single motor transported the GeNTOP is $p_1 = (1-p_m)(1-\ln(1-p_m))$ not accounting for that a motor, bound opposite to another one, may not be able to interact simultaneously. For single-molecule experiments, the pipetted kinesin-to-GeNTOP ratio was about 20 corresponding to a motile fraction of $p_m \lesssim 30\,\%$ implying single-molecule conditions with at least 95 % confidence. To measure speed and run length of single kinesin motors on microtubules in the absence of loads, we used another custom-built optical tweezers setup combined with interference reflection microscopy (IRM) (*61*). Motor-coated GeNTOPs were trapped and placed on a microtubule. If the GeNTOP showed motility, the trap was turned off and IRM images were acquired at a rate of 7 frames/s. Based on kymographs, the mean motor speed and run length was $0.72 \pm 0.05\,\mu$m/s and $1.1 \pm 0.4\,\mu$m ($N = 12$), respectively, consistent with literature values (*62–64*).

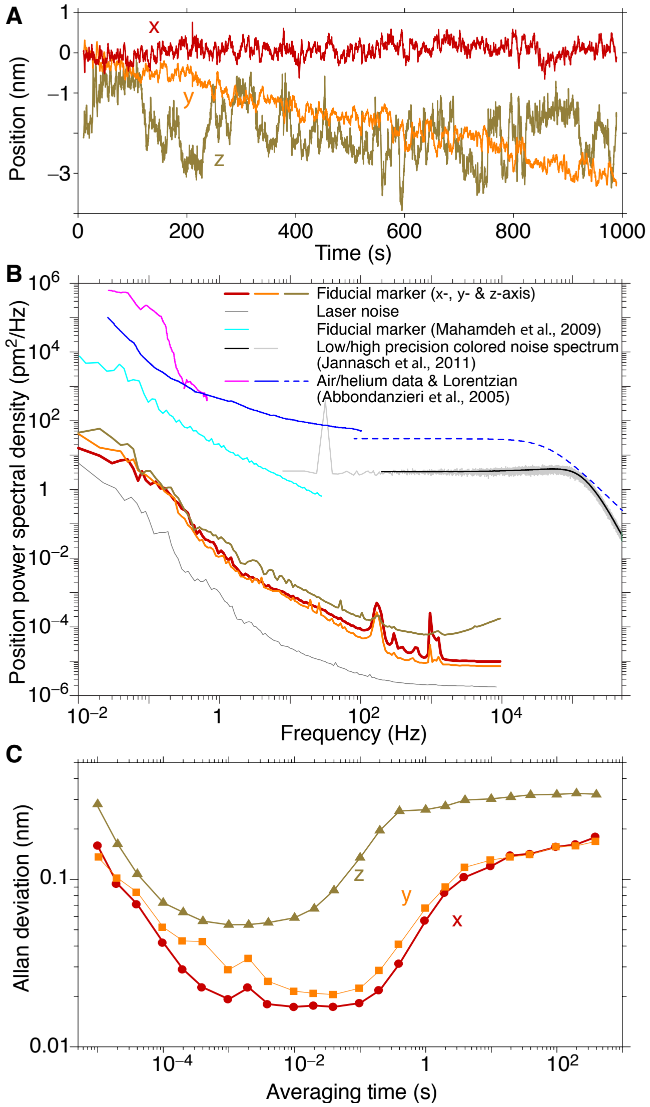

**Fig. S1. Ultrastable optical tweezers. (A)**, Position of a fiducial marker as a function of time (100 kHz data blocked to 10 Hz bandwidth). **(B)**, Position power spectral density recorded for a fiducial marker in comparison to the stability of the benchmark setup (Abbondanzieri *et al.*, 2005 (*65*)) and previously recorded data (Mahamdeh *et al.*, 2009 (*49*) and Jannasch *et al.*, 2011 (*54*)). **(C)**, Allan deviation as a function of lag time for the same data. Note that the setup was moved from a third-floor laboratory at the TU Dresden, Germany, where previous data (*49, 54*) was recorded, to a basement room at the University of Tübingen, Germany, with excellent vibration and sound isolation and temperature stability (*66*).

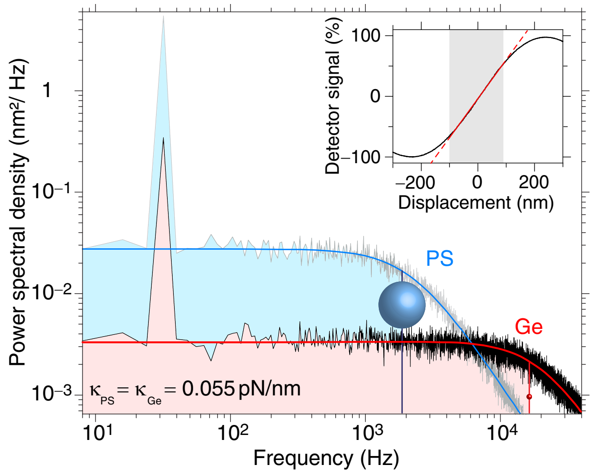

**Fig. S2. Spatiotemporal precision of optically trapped germanium nanospheres (GeNTOPs).** Power spectral density (average of 40 individual power spectra) of GeNTOP (70-nm diameter, germanium (Ge), black line) and microsphere motion (0.59-μm diameter, polystyrene (PS), grey line) trapped in water. Spectra feature a calibration peak at 32 Hz (red and blue lines, fit to theory (*50*), see methods). Corner frequencies $f_c$ are indicated by vertical lines through schematic, proportionally scaled spheres. The corner frequency serves as a measure for the available measurement bandwidth (shaded areas). Inset: lateral detector response of a surface-immobilized GeNTOP as a function of displacement relative to the trap centre (black line, linear fit red line). Because of the fluctuation-dissipation theorem, the area underneath the power spectra of the GeNTOP and microsphere motion is the same. However, power is distributed differently across the frequency space with a higher corner frequency and lower positional noise level at low frequencies for the GeNTOPs compared to the microsphere allowing for an improved spatiotemporal resolution.

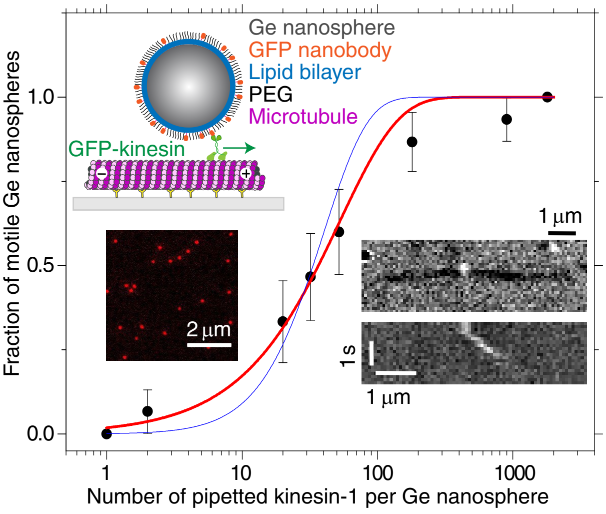

**Fig. S3. Single kinesins transported lipid-bilayer-coated GeNTOPs.** Fraction of motile GeNTOPs as a function of kinesin-to-GeNTOP ratio. Data (black circles, 40 tested nanospheres per condition) with Poisson statistics fit (transport by at least one (red line) or at least two (blue line) motors, see methods). Inset: Schematic of a kinesin motor transporting a functionalized GeNTOP along a microtubule drawn roughly to scale (top left). Confocal image of lipid bilayer-coated GeNTOPs with a membrane dye confirmed the presence of the lipid bilayer (left). Interference reflection microscopy image and kymograph (right) of a single kinesin-transported GeNTOP placed on a microtubule with the optical tweezers (bright and dark contrast, respectively).

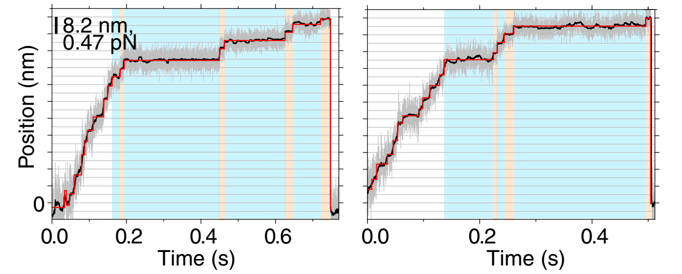

**Fig. S4. Exemplary kinesin traces at physiological ATP concentrations.** Time traces for a single-kinesin powered GeNTOP (100 kHz bandwidth, grey trace; filtered data, ≈100 Hz, black trace; detected steps, red line; see methods). Long and short dwell times $\tau_1$ and $\tau_2$ are blue and orange shaded, respectively.

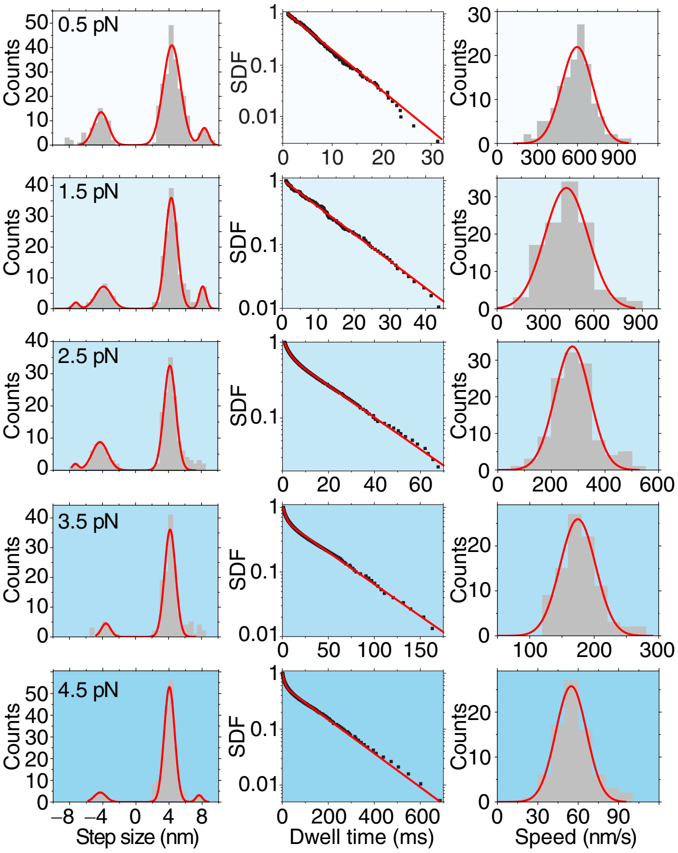

**Fig. S5. Step size, dwell time, and speed distributions versus force at physiological ATP concentrations.** Step size histograms with multi-Gaussian fit (left column), survival distribution functions (SDFs) of dwell times with fits of single or sum of two exponentials (middle column) and speed histograms with Gaussian fit (right column) for forces range with centres from 0.5 pN to 4.5 pN (top to bottom). See table S1 for fit results.

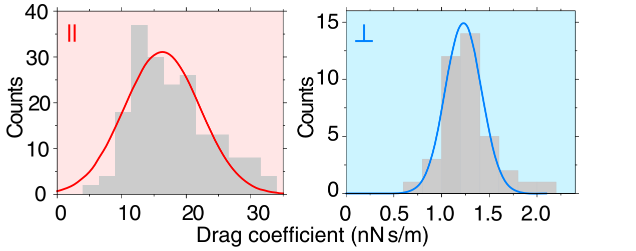

**Fig. S6. Frictional drag coefficient based on fast backward motion.** Histograms of the frictional drag coefficient measured parallel ($\gamma_{||}$, left, red shaded) and perpendicular ($\gamma_{\perp}$, right, blue shaded) to the microtubule axis with Gaussian fits (red and blue lines). The frictional drag coefficient for the two directions was calculated according to $\gamma = \tau\kappa$ using the measured values for the relaxation time and trap stiffness (see Fig. 3 and methods). The resulting values for $\gamma_{||}$ and $\gamma_{\perp}$ are 16.0 ± 0.8 nN s/m ($N = 149$) and 1.24 ± 0.07 nN s/m ($N = 50$), respectively. The latter frictional drag coefficient $\gamma_{\perp}$ was larger than the hydrodynamic (viscous) drag coefficient expected from Stokes drag and the measured GeNTOP size. The ratio between the measured coefficient $\gamma_{\perp}$ and the calculated Stokes drag coefficient is about 1.9. This increase is due to the surface proximity (*43*). Based on Faxén's law, this ratio is consistent with the GeNTOP being 10 nm away from the surface.

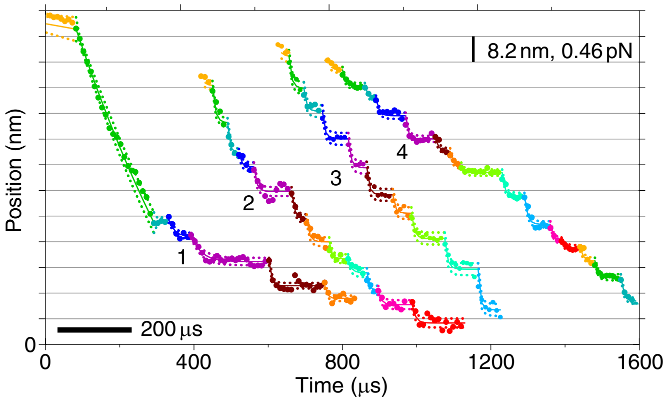

**Fig. S7. Exemplary kinesin traces of fast, biased sliding motion in the weakly bound diffusive state.** Coloured sections correspond to detected states fitted with a single exponential relaxation using the "Steppi" algorithm (*59*). Traces 1 and 2 had no constraints resulting in a step size of 7.7 ± 0.1 nm and exponential relaxation time of 27 ± 3 µs ($N = 20$, excluding the first step of Trace 1). For Traces 3 and 4, the relaxation time constant was fixed to 25 µs (see methods). The force scale bar is based on the average trap stiffness for the four traces.

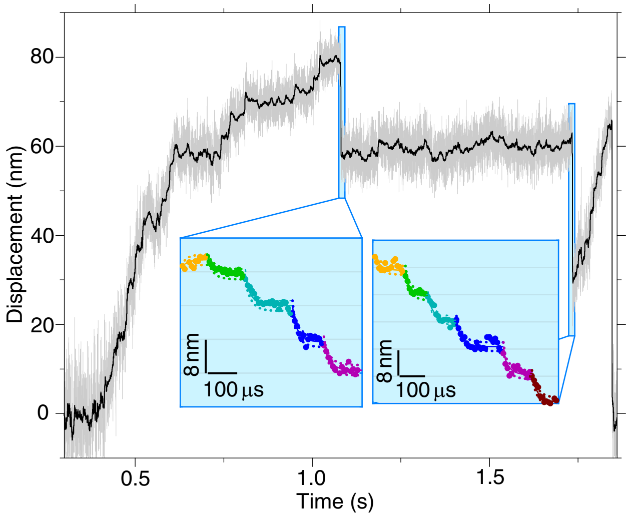

**Fig. S8. Exemplary kinesin trace with short slip events.** Time traces for a single-kinesin powered GeNTOP (100 kHz bandwidth, grey trace; filtered data, ≈100 Hz, black trace; see methods). Insets: Magnified view of short backward slips. Coloured sections correspond to detected states fitted with a single exponential relaxation using the "Steppi" algorithm (*59*).

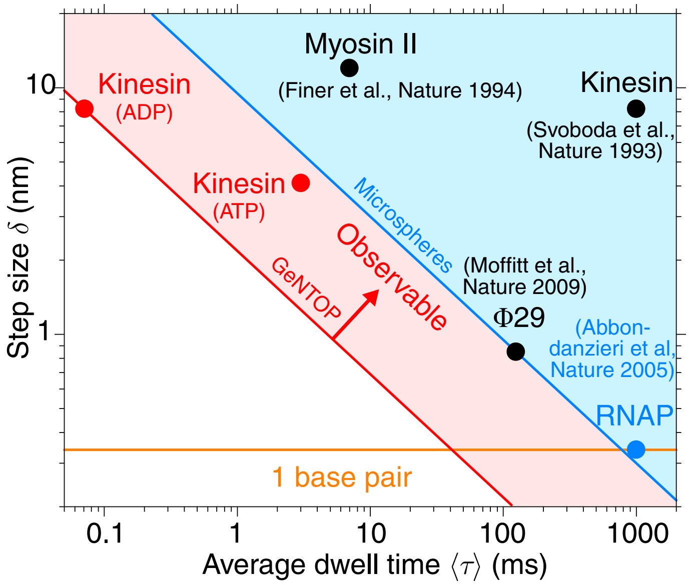

**Fig. S9. Optical tweezers spatiotemporal resolution of molecular machines.** Step size versus dwell time for various molecular machines (*65, 67–69*) in comparison to this work (red circles, 4-nm directed substeps (0–1 pN data point at 10 µM ATP of table S2), fast 8-nm sliding steps (ADP state, Fig. 3A)). Blue and red line indicate previous, microsphere benchmark (*65*) and current GeNTOP spatiotemporal resolution (this work), respectively, according to the relation $\delta\sqrt{\langle\tau\rangle}$ (*70*). The half space above the lines is observable.

**Table S1. Step size, dwell time and speed versus force at 1 mM ATP.**

| $F$ | $\delta_+$ (nm) | $p$ | $\delta_-$ (nm) | $p$ | $\tau_1$ (ms) | $p$ | $\tau_2$ (ms) | $p$ | $v$ (µm/s) |
|---|---|---|---|---|---|---|---|---|---|
| 0–1 | $4.34 \pm 0.06$ (217) | 72 | $4.2 \pm 0.2$ (65) | 22 | $5.5 \pm 0.2$ (306) | 120 | – | – | $0.60 \pm 0.01$ (134) |
| | $8.2 \pm 0.6$ (19) | 6 | | | | | | | |
| 1–2 | $4.25 \pm 0.04$ (140) | 72 | $4.0 \pm 0.2$ (34) | 18 | $10.3 \pm 0.8$ (193) | 103 | – | – | $0.43 \pm 0.01$ (110) |
| | $7.8 \pm 0.2$ (15) | 8 | $8.2 \pm 0.6$ (4) | 2 | | | | | |
| 2–3 | $4.19 \pm 0.04$ (127) | 71 | $4.1 \pm 0.2$ (42) | 25 | $20.6 \pm 0.1$ (100) | 71 | $2.1 \pm 0.7$ (78) | 50 | $0.28 \pm 0.01$ (117) |
| | $7.3 \pm 0.5$ (4) | 2 | $7.3 \pm 0.5$ (5) | 2 | | | | | |
| 3–4 | $4.18 \pm 0.05$ (127) | 85 | $3.9 \pm 0.4$ (16) | 10 | $44.7 \pm 0.4$ (84) | 63 | $3.3 \pm 0.7$ (66) | 49 | $0.18 \pm 0.01$ (112) |
| | $7.8 \pm 0.2$ (7) | 5 | | | | | | | |
| ≥4 | $4.06 \pm 0.03$ (168) | 89 | $4.3 \pm 0.4$ (16) | 8 | $144 \pm 2$ (104) | 60 | $8.8 \pm 0.2$ (86) | 47 | $0.055 \pm 0.001$ (92) |
| | $7.6 \pm 0.4$ (6) | 3 | | | | | | | |

$F$ (pN): force, $\delta_{+/-}$: forward/backward step size—Gaussian centre ± fit error ($N$ based on area underneath Gaussian normalized by total number of steps), $p$ (%): relative percentage, $\tau$: dwell time based on survival function fit ($N$ according to relative amount), and $v$: speed—mean ± standard error ($N$: number of trace segments fitted). All fits to data of fig. S5. Note that only few data points correspond to forces larger than 5 pN. Also note that $p$-values for dwell times directly reflect the fitted amplitude that may add up to more than 100 % indicating that some of the expected very short steps were missed. Errors on all percentages are less than 1 %.

**Table S2. Step size, dwell time and speed versus force at 10 µM ATP.**

| $F$ | $\delta_+$ (nm) | $p$ | $\delta_-$ (nm) | $p$ | $\tau_1$ (ms) | $p$ | $\tau_2$ (ms) | $p$ | $v$ (µm/s) |
|---|---|---|---|---|---|---|---|---|---|
| 0–1 | $4.13 \pm 0.06$ (53) | 67 | $4.2 \pm 0.2$ (15) | 20 | $30.2 \pm 0.6$ (44) | 65 | $3.1 \pm 0.2$ (34) | 47 | $0.23 \pm 0.01$ (12) |
| | $7.2 \pm 0.2$ (10) | 13 | | | | | | | |
| 1–2 | $4.15 \pm 0.03$ (53) | 69 | $4.5 \pm 0.4$ (9) | 11 | $37.8 \pm 0.7$ (42) | 61 | $3.3 \pm 0.1$ (34) | 49 | $0.16 \pm 0.01$ (17) |
| | $7.6 \pm 0.36$ (15) | 20 | | | | | | | |
| 2–3 | $4.05 \pm 0.03$ (59) | 78 | $4.5 \pm 0.5$ (7) | 9 | $54.3 \pm 0.1$ (40) | 58 | $3.8 \pm 0.1$ (37) | 52 | $0.110 \pm 0.007$ (13) |
| | $7.5 \pm 0.17$ (10) | 13 | | | | | | | |
| 3–4 | $4.17 \pm 0.05$ (42) | 70 | $4.3 \pm 0.2$ (6) | 10 | $131 \pm 3$ (33) | 57 | $6.8 \pm 0.3$ (27) | 45 | $0.056 \pm 0.005$ (15) |
| | $7.84 \pm 0.36$ (12) | 20 | | | | | | | |
| ≥4 | $4.12 \pm 0.04$ (77) | 85 | $3.9 \pm 0.2$ (6) | 7 | $245 \pm 5$ (41) | 56 | $8.0 \pm 0.3$ (40) | 45 | $0.032 \pm 0.004$ (15) |
| | $8.12 \pm 0.13$ (7) | 8 | | | | | | | |

$F$ (pN): force, $\delta_{+/-}$: forward/backward step size—Gaussian centre ± fit error ($N$ based on area underneath Gaussian normalized by total number of steps), $p$ (%): relative percentage, $\tau$: dwell time based on survival function fit ($N$ according to relative amount), and $v$: speed—mean ± standard error ($N$: number of trace segments fitted). Note that only few data points correspond to forces larger than 5 pN. Also note that $p$-values for dwell times directly reflect the fitted amplitude that may add up to more than 100 % indicating that some of the expected very short steps were missed. Errors on all percentages are less than 1 %.

### References and Notes

37. Y. J. Guo, *et al.*, *Chem. Asian J.* **9**, 2272 (2014).
38. L. Ma, Y. Cai, Y. Li, J. Jiao, Z. Wu, *elife* **6**, 1 (2017).
39. P. B. Santhosh, N. Thomas, S. Sudhakar, A. Chadha, E. Mani, *Phys. Chem. Chem. Phys.* **19**, 18494 (2017).
40. I. Brouwer, *et al.*, *Nat. Commun.* **6**, 1 (2015).
41. M. Bugiel, *et al.*, *J. Biol. Methods* **2**, 30 (2015).
42. U. Rothbauer, *et al.*, *Mol. Cell Proteomics.* **7**, 282 (2008).
43. E. Schäffer, S. F. Nørrelykke, J. Howard, *Langmuir* **23**, 3654 (2007).
44. V. Bormuth, V. Varga, J. Howard, E. Schäffer, *Science* **325**, 870 (2009).
45. A. Ramaiya, B. Roy, M. Bugiel, E. Schäffer, *Proc. Natl. Acad. Sci. USA* **114**, 10894 (2017).
46. M. Bugiel, E. Böhl, E. Schäffer, *Biophys. J.* **108**, 2019 (2015).
47. M. Bugiel, A. Jannasch, E. Schäffer, *Optical Tweezers: Methods and Protocols*, A. Gennerich, ed. (Humana Press, 2016), chap. 5, pp. 109–136.
48. M. Mahamdeh, Campos, C.P, E. Schäffer, *Opt. Express* **19**, 11759 (2011).
49. M. Mahamdeh, E. Schäffer, *Opt. Express* **17**, 17190 (2009).
50. S. F. Tolic-Nørrelykke, E. Schäffer, J. Howard, F. S. Pavone, F. Jülicher, *Rev. Sci. Instrum.* **77**, 103101 (2006).
51. T. A. Nieminen, *et al.*, *J. Opt. A: Pure Appl. Opt.* **9**, 196 (2007).
52. V. Bormuth, *et al.*, *Opt. Express* **16**, 13831 (2008).
53. https://www.filmetrics.com/refractive-index-database/Ge/Germanium. Accessed 23.07.2020.
54. A. Jannasch, M. Mahamdeh, E. Schäffer, *Phys. Rev. Lett.* **107**, 228301 (2011).
55. E. J. G. Peterman, F. Gittes, C. F. Schmidt, *Biophys. J.* **84**, 1308 (2003).
56. T. J. Jachowski, Stepfinder: A Python package to find steps in one dimensional data with low SNR., GitHub repository: https://github.com/tobiasjj/stepfinder (2019).
57. S. H. Chung, R. A. Kennedy, *J. Neurosci. Methods* **40**, 71 (1991).
58. D. A. Smith, *Philos. Trans. R. Soc. B* **353**, 1969 (1998).
59. P. A. Wiggins, *Biophys. J.* **109**, 346 (2015).
60. K. Svoboda, S. M. Block, *Cell* **77**, 773 (1994).
61. S. Simmert, M. Abdosamadi, G. Hermsdorf, E. Schäffer, *Opt. Express* **26**, 1437 (2018).
62. S. Block, L. Goldstein, B. Schnapp, *Nature* **348**, 348 (1990).
63. D. L. Coy, M. Wagenbach, J. Howard, *J. Biol. Chem.* **274**, 3667 (1999).
64. D. Cai, K. J. Verhey, E. Meyhöfer, *Biophys. J.* **92**, 4137 (2007).
65. E. A. Abbondanzieri, W. J. Greenleaf, J. W. Shaevitz, R. Landick, S. M. Block, *Nature* **438**, 460 (2005).
66. G. L. Hermsdorf, S. A. Szilagyi, S. Rösch, E. Schäffer, *Rev. Sci. Instrum.* **90**, 015113 (2019).
67. K. Svoboda, C. F. Schmidt, B. J. Schnapp, S. M. Block, *Nature* **365**, 721 (1993).
68. J. T. Finer, R. M. Simmons, J. A. Spudich, *Nature* **368**, 113 (1994).
69. J. R. Moffitt, *et al.*, *Nature* **457**, 446 (2009).
70. J. R. Moffitt, Y. R. Chemla, S. B. Smith, C. Bustamante, *Annu. Rev. Biochem.* **77**, 205 (2008).
